# TCRβ Sequencing Reveals Spatial and Temporal Evolution of Clonal CD4 T cell Responses in a Breach of Tolerance Model of Inflammatory Arthritis

**DOI:** 10.1101/2020.11.15.383539

**Authors:** Shaima Al-Khabouri, Robert A. Benson, Catriona T. Prendergast, Joshua I. Gray, Thomas D Otto, James M Brewer, Paul Garside

## Abstract

Effective tolerogenic intervention in Rheumatoid Arthritis (RA) will rely upon understanding the evolution of articular antigen specific CD4 T cell responses. TCR clonality of endogenous CD4 T cell infiltrates in early inflammatory arthritis was assessed to monitor evolution of the TCR repertoire in the inflamed joint and associated lymph node (LN). Mouse models of antigen-induced breach of self-tolerance and chronic polyarthritis were used to recapitulate early and late phases of RA. The infiltrating endogenous, antigen experienced CD4 T cells in inflamed joints and LNs were analysed using flow cytometry and TCRβ sequencing. TCR repertoires from inflamed late phase LNs displayed increased clonality and diversity compared to early phase LNs, while inflamed joints remained similar with time. Repertoires from late phase LNs accumulated clones with a diverse range of TRBV genes, while inflamed joints at both phases contained clones expressing similar TRBV genes. Repertoires from LNs and joints at the late phase displayed reduced CDR3β sequence overlap compared to the early disease phase, however the most abundant clones in LNs accumulate in the joint at the later phase. The results indicate CD4 T cell repertoire clonality and diversity broadens with progression of inflammatory arthritis and is first reflected in LNs before mirroring in the joint. These observations imply that antigen specific tolerogenic therapies could be more effective if targeted at earlier phases of disease when CD4 T cell clonality is least diverse.

## INTRODUCTION

Rheumatoid arthritis (RA) is a chronic inflammatory autoimmune disease characterised by synovial inflammation and cartilage and bone erosion, causing progressive loss of joint function (1). Antigen presentation and CD4 T activation mechanisms have been shown to play a role in the pathogenesis of RA. This is evidenced by genetic association studies in RA patients showing strong associations of HLA-DRB alleles, T cell activation associated genes such as *PTPN22* and *CTLA-4*, and loci associated with signal transduction such as *STAT4* (2–7). Moreover, the successes of therapeutics in modulating T cell activation, such as the use of Abatacept, reinforce the importance of CD4 T cells in propagating the disease (8). As such, there has been increasing interest in targeting pathogenic autoreactive CD4 T cells using antigen specific tolerogenic therapies as this line of therapy aims to re-establish tolerance and provide long-lasting remission, while retaining protective immunity against pathogens. However, targeting autoreactive CD4 T cells has proven difficult in RA, due to the broad range of antigens implicated in disease, lack of a clear cellular hierarchy of disease drivers, and also patient to patient variation (9–13). Moreover, it is unknown at what stages of the disease these autoreactive responses develop nor the location in which these responses are primed, thus hampering the development of effective tolerogenic therapies. Antigen specific responses can be identified by detecting expanded clonal T cell populations. Indeed, oligoclonal CD4 T cell responses and skewed TCR repertoires have been reported in arthritic joints (14–17). More recently, analysis of CD4 T cell behaviour was shown to reflect antigen recognition in inflamed joints at the early stages of disease (18), suggesting that antigen specific CD4 T cell responses may perpetuate and drive progression of RA. However, the development of antigen specific responses and their evolution with the progression of RA is poorly understood. Investigating the evolution TCR clonality of antigen experienced CD4 T cells in RA will provide insight on how antigen specific responses evolve with disease progression. This will inform the development of more effective tolerogenic therapies by indicating the range of clones that would need to be targeted and the disease stage at which these therapies are most likely to be effective. In this study, we employed breach of self-tolerance models of antigen induced inflammatory arthritis (19,20) and chronic polyarthritis (21) in which autoreactive responses develop. These models also recapitulate the early and later stages of the disease – hereafter named the early and late phases respectively. TCRβ sequencing was used to monitor the evolution of antigen specific endogenous CD4 T cell responses in inflamed joints and their associated draining lymph nodes.

## METHODS

### Animals

Male and female C57BL/6 mice aged 7-12 weeks were purchased from Envigo (UK), and used as adoptive transfer recipients. OT-II TCR transgenic mice (22) were bred inhouse. All mice were maintained at the University of Glasgow’s central research facility and housed under standard housing conditions specified by the UK Home Office.

### Induction of inflammatory arthritis

“Early” antigen induced inflammatory arthritis was conducted as previously described (19,20). Briefly, 2-3×10^6^ Th1 polarised OT-II transgenic T cells were transferred i.v. into C57BL/6 recipients. Recipients were immunised with 100µL of 1µg/mL of OVA protein (Worthington Biochemicals) in Freund’s complete adjuvant (CFA) (Sigma-Aldrich) subcutaneously after 24 hours. Ten days later, mice were given a periarticular injection in footpads with 50µL of 100µg of heat aggregated OVA (HAO) or PBS as a control. Mice were sacrificed four days later and cells were isolated from joint tissue and draining popliteal lymph nodes (pLN). The “late” model antigen induced inflammatory arthritis was modified from a model of chronic polyarthritis(21). 30 days after HAO administration described above, mice were given a second periarticular injection of 100µg of HAO in incomplete Freund’s adjuvant (IFA) (Sigma-Aldrich) or IFA alone as a control. Mice were sacrificed 17 days later and cells were isolated from joint tissues and pLNs.

### Isolation of cells from tissues

Cells from mouse joints and pLNs were isolated as previously described (18). Briefly, ankle joints were teased apart and shaken at 110 rpm at 37°C for 25 minutes with 2.68mg/mL collagenase D (Roche) in RPMI 1640 (Gibco, Thermo Fisher Scientific). Joint tissue was then homogenised using a gentleMACS Dissociator (Miltenyi Biotech) then strained to obtain single cell suspensions. PLNs were forced through Nitex mesh (Cadisch Precision Meshes) to obtain single cell suspensions. Samples were washed and stained for flow cytometry. Joint and pLN samples were analysed separately. Joint samples and pLN samples were pooled per mouse.

### Flow cytometry

Cells were stained for flow cytometry as described previously (18). The following antibodies were used: anti-CD4 (clone GK1.5), anti-CD44 (clone IM7), anti-CD45 (clone 30-F11), anti-CD45.1 (clone A20). Data was analysed using FlowJo v10 software (Treestar, Oregon USA).

### TCRβ Sequencing

The data for this study have been deposited in the European Nucleotide Archive (ENA) at EMBL-EBI under the accession number PRJEB40509 (www.ebi.ac.uk/ena/browser/view/PRJEB40509). CDR3β sequencing was performed by iRepertoire Inc. (Huntsville, AL, USA) on total RNA collected from sorted antigen experienced endogenous CD4 T cells (CD4+, C45.1-, CD44hi). CD4+, CD45.1-, CD44hi were sorted from joint and pLN samples using the BD Aria III FACS sorter (BD Biosciences). Samples were collected separately from individual mice in lysis buffer (Purelink RNA micro kit, Thermofisher) + 1% β- mercaptoethanol (Sigma-Aldrich) and lysed using 29G insulin syringes (VWR). Lysed cells were then stored at −80°C prior to RNA purification. Total RNA was purified using the Purelink RNA micro kit (Thermo Fisher Scientific) as per manufacturer’s instructions and quality assessed by measuring A260/280 using a Nanodrop 1000 spectrophotometer (Thermo Fisher Scientific). Barcoded libraries were prepared per sample using barcoded primers covering the V-J TCR region, then amplified, pooled, and sequenced using the Illumina MiSeq platform covering 150 paired end reads (PER). Basic data analysis was performed by iRepertoire. Sequencing data was also prepared for analysis using MiXCR developed by Bolotin et. al (23) and analysed using tcR developed by Nazarov et. al (24) in the R statistical software frame work (R version 3.4.3).

### Calculation of the D50 diversity index (DI)

Diversity was calculated using iRepertoire’s data analysis guide. For samples where the number of unique CDR3β sequences ≥10,000: DI = rank of unique CDR3β sequence where 50% of the top 10,000 total reads falls x 100 / 10,000

For samples where the number of unique CDR3β sequences <10,000: DI = rank of unique CDR3β sequence where 50% of the total reads falls x 100 / no. of unique CDR3β sequences

### Calculation of repertoire overlap

The number of shared CDR3β amino acid sequences is represented as a normalised overlap index taking into consideration the size of the repertoires being compared. The normalised overlap index was calculated using the tcR R package (24) using the following calculation: Normalised overlap index = No. of exact overlapping CDR3β amino acid sequences/ total reads repertoire 1 x total number of reads repertoire 2

### Statistics

Data is shown as mean ± SD. Specific test and significance levels are stated in respective figure legends. Statistical analyses were performed using GraphPad Prism version 7 (GraphPad Inc, CA, USA) or using the R statistical software framework (R version 3.4.3).

## RESULTS

### Induction of inflammatory arthritis results in local accumulation of endogenous antigen experienced CD4 T cells at both the early and late phases

Induction of the breach of tolerance model of inflammatory arthritis requires the transfer of Th1 polarised ovalbumin (OVA) specific OT-II cells and subsequent immunisations with OVA and heat aggregated OVA (HAO) as outlined in **figure 1 A**. This results in the influx of the transferred OT-II cells and endogenous CD4 T cells of varying antigen specificities, indicated by the range of expressed TCRβ variable genes (TRBV) (18). Assessing the evolution of TCR clonality of the endogenous CD4 T cell population requires the accurate identification of these cells in joints and popliteal lymph nodes (pLN), being distinguishable from the transferred OT-IIs by expression of the congenic marker CD45.1 **(supplementary figures 1 and 2)**. Endogenous CD4 T cells were present at a significantly higher number than the transferred OT-II cells in inflamed pLNs and joints at both the early and late phases **(figure 1 B)**. A significant number of endogenous CD4 T cells at the early phase also displayed an antigen experienced phenotype, indicated by increased CD44 expression, after induction of inflammatory arthritis with HAO **(figure 1 C)**, and were previously shown to produce TNFα and IFNγ following ex-vivo stimulation with PMA/ionomycin (18). The number of endogenous CD4, CD44hi cells was also increased at the late phase in inflamed joints **(figure 1 C)**.

**Figure 1.**
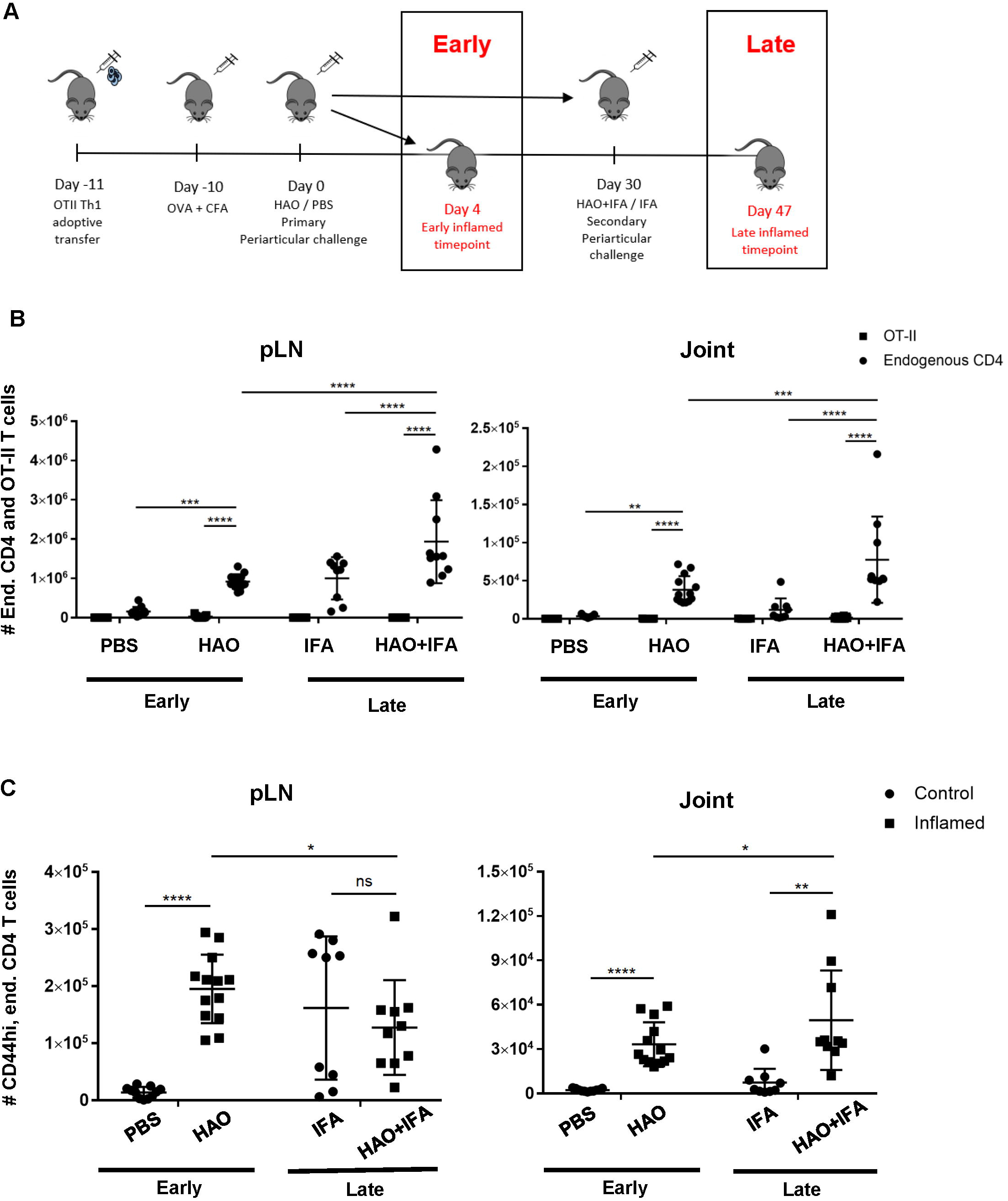
Outline of model phases and number of endogenous, OT-II, and endogenous antigen experienced CD4 T cells at both phases. **A)** Illustration of early and late phases of inflammatory arthritis. **B)** Number of endogenous CD4 T cells (CD4+, CD45+, CD45.1-) and OT-II T cells (CD4+, CD45+, CD45.1+) from pLNs and joints of mice undergoing inflammatory arthritis at the early and late phases and respective controls. **C)** Number of antigen experienced endogenous CD4 T cells (CD4+, CD45+, CD45.1-, CD44hi) from pLNs and joints of mice undergoing inflammatory arthritis at the early and late phases and respective controls. Data is representative of mean ± SD with each point representing individual experimental mice. Data represents three independent experiments for the early phase and two independent experiments for the late phase with n=5 for each experiment. Groups were compared using 2-way ANOVA and Student’s t-tests. Stars represent the following p-values: *<0.05, ** <0.01; ***<0.001; **** <0.0001; ns: not significant. HAO, heat-aggregated OVA; IFA, incomplete Freund’s adjuvant.

**Figure 2.**
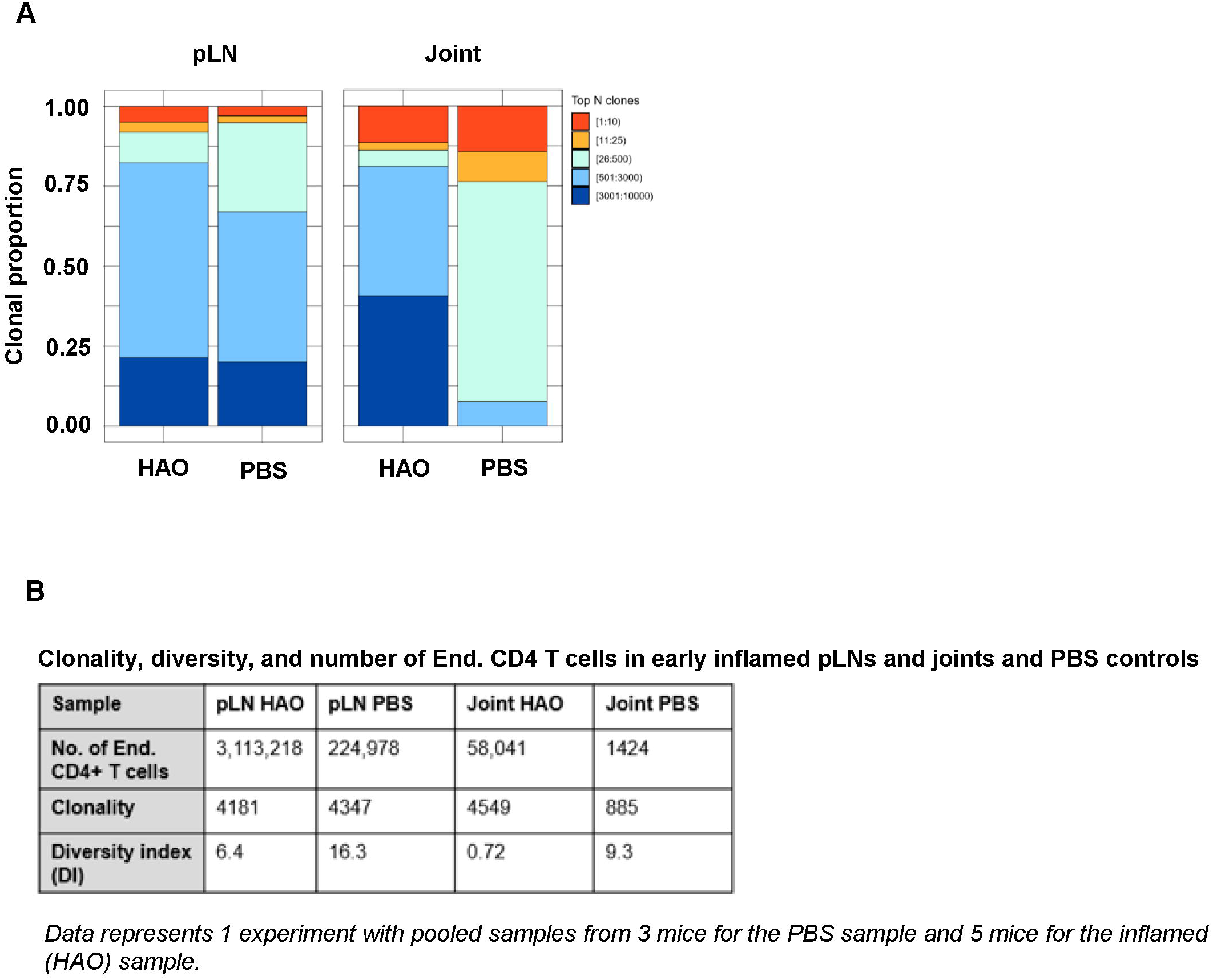
Characterisation of antigen experienced endogenous CD4 T cell repertoires from inflamed pLNs and joints and PBS controls at the early phase. Proportion of the top 10 most frequently occurring clones, followed by the next 25, 500, 3000, 10,000, and 100,000 most frequent clones contributing to the endogenous antigen experienced CD4+ T cell repertoire in pooled inflamed pLNs and joint samples at the early phase (HAO) and pooled PBS controls. Data represents 1 experiment with pooled samples from 3 mice for the PBS sample and 5 mice for the inflamed (HAO) sample.

### Accumulation of expanded clones and observed clonality and diversity of CD4 T cell repertoires is antigen driven

To investigate the evolution of CD4 TCR clonality in this model of inflammatory arthritis, endogenous antigen experienced CD4 T cells i.e. excluding the transferred OT-II cells, were sorted from pLNs and joints of individual mice at both early and late phases. Total RNA was then isolated from the endogenous CD4 T cells and used to sequence the CDR3β region of the TCR, the most variable region of the TCR critical in recognition of specific peptide sequences (25,26). Repertoires were then characterised by examining the number of unique CDR3β sequences and evaluating the degree expanded clones contributed to the overall antigen experienced repertoire. From this, the clonality and diversity of the CD4 T cell repertoire can be determined. In this study, clonality is defined by the number of unique CDR3β sequences found in the population, and diversity – represented by the Diversity Index (DI) (see Methods) □ represents the relative contribution of each unique CDR3β sequence, or clone, to the population. Thus, a repertoire with a larger number unique CDR3β sequences, and therefore clones, would be described to have high clonality. This repertoire would also display high diversity and have a high DI value if the number of cells contributing to different clonal populations are more evenly distributed. Conversely, the repertoire would be described to have low diversity and have a low DI value if the repertoire is mainly comprised of a few clones present in high frequencies relative to the total number of clones. Due to the low recovery of antigen experienced endogenous CD4 T cells isolated from control pLNs and joints at the early phase, cells from control pLNs or joints needed to be pooled prior to sequencing. Pooling samples changes the composition of the repertoire and affects T cell repertoire diversity, so comparisons of control CD4 T cell populations were made against pooled data from inflamed joint and pLN samples at the early phase **(figures 2 A and B)**. Clonality of the endogenous CD4 T cell repertoire from inflamed pLNs at the early phase and controls were comparable (4181 unique CDR3β sequences in the inflamed pLN repertoire vs 4347 in the PBS pLN repertoire). However, the repertoire from the inflamed pLN showed a reduced diversity compared with the control pLN sample (6.4 vs 16.3 respectively). Endogenous CD4 T cell repertoires from inflamed joints displayed higher clonality than controls (4549 vs 885). Despite this, the diversity of the repertoire in the inflamed joint was relatively low compared to the PBS joint (0.72 vs 9.3 respectively). These data therefore demonstrate that accumulation of clones can be detected at the early phases of the disease in both inflamed pLNs and joints.

Evidence for the accumulation of antigen experienced T cells in the absence of their cognate antigen has been demonstrated in this model (18) and in RA patients (27–29) which may be a result of a chronic inflammatory environment (30,31). To address whether the accumulation of clones is antigen driven rather than a result of non-antigen induced inflammation, the antigen experienced repertoires of pLNs and joints at the late phase were compared to repertoires isolated from mice challenged with IFA alone **(figure 3 A)**, where inflammation is induced without the presence of antigen. No significant difference was found between the contribution of the top 10 most expanded clones to the repertoires of inflamed pLNs at the late phase and IFA controls (data not shown) nor when comparing the contribution of clones occurring at the lowest frequency **(figure 3 B)**. The number of clones occurring at a frequency of 2 or more were also compared between inflamed and IFA pLN samples as this was the median frequency of clones found in the PBS pLN sample (data not shown). Thus, clonal populations found at higher than this frequency are assumed to have accumulated due to the presence of antigen. No clonal accumulation was detected between inflamed pLNs at the late phase and IFA controls (*p value 0*.*32 Welch two sample t-test*). Moreover, repertoires from inflamed pLNs at the late phase and IFA displayed comparable levels of clonality and diversity **(figures 3 C and D)**.

**Figure 3.**
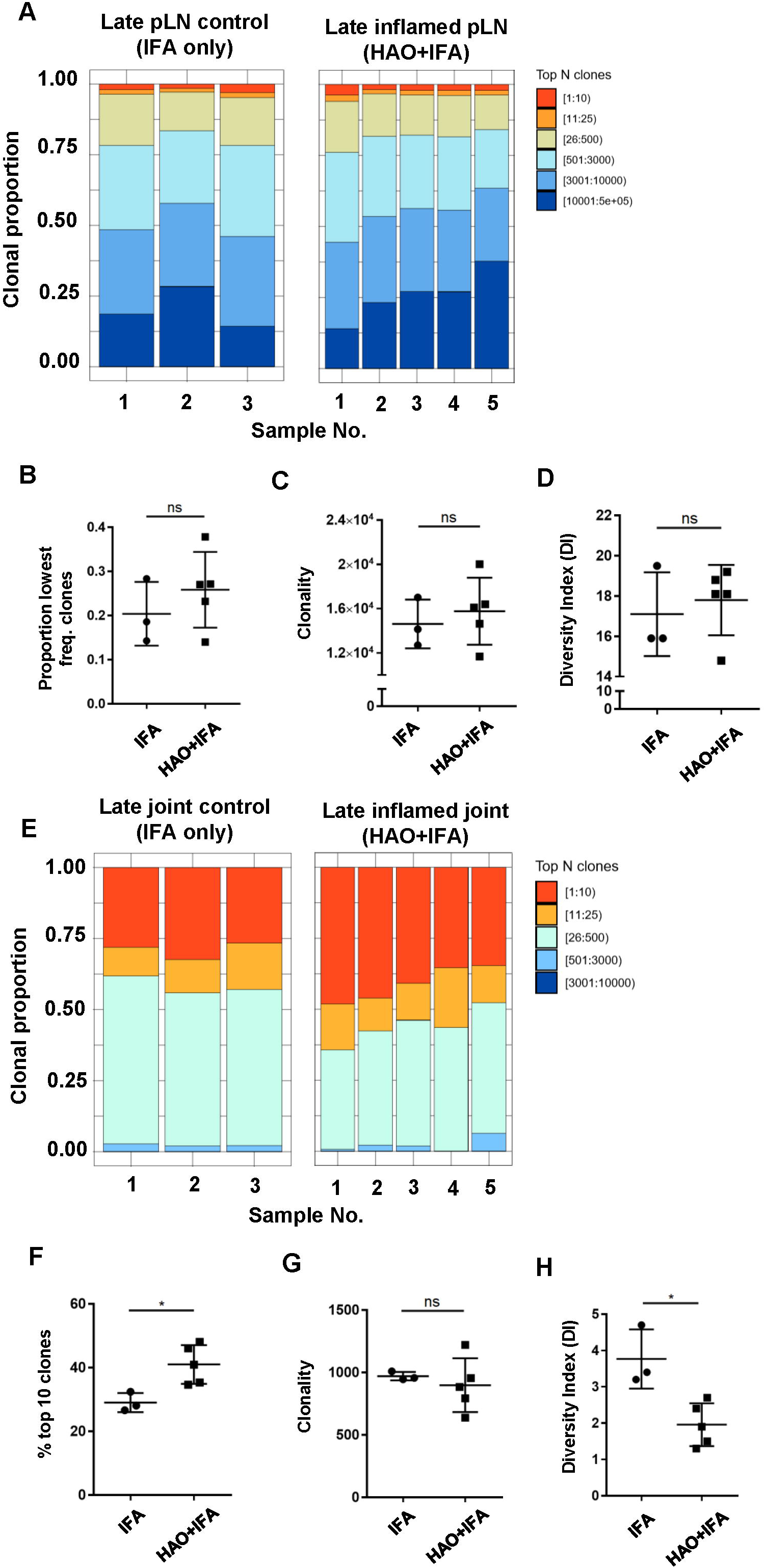
Characterisation of antigen experienced endogenous CD4 T cell repertoires from inflamed pLNs and joints and IFA controls at the late phase. Proportion of the top 10 most frequently occurring clones, followed by the next 25, 500, 3000, 10,000, and 100,000 most frequent clones contributing to the endogenous antigen experienced CD4+ T cell repertoire in **A)** inflamed pLNs, and **E)** inflamed joints at the late phase (HAO+IFA) and IFA only controls. **B)** Proportion of the least frequently occurring clones in inflamed pLNs at the late phase and IFA controls. **F)** Percentage contribution of the top 10 most frequently occurring clones in inflamed joints at the late phase and IFA controls. **C and G** clonality, represented as the number of unique CDR3β DNA sequences, in **C)** inflamed pLNs and IFA controls, and **G)** inflamed joints and IFA controls at the late phase. **D and H** diversity indices of **D)** inflamed pLNs and IFA controls, and **H)** inflamed joints and IFA controls at the late phase. Data is representative of mean ± SD with each point representing individual experimental mice. Data represents 1 experiment with n=5 for the inflamed group and n=3 for the control group. Groups were compared using unpaired Student’s t-test. Stars represent the following p values: * <0.05; ns: not significant.

In late phase arthritic joints, the top 10 highly expanded clones constituted approximately 41% of the overall repertoire compared to joint IFA controls, where the top clones only comprised an average of 29% of the entire repertoire **(figures 3 E and F)**. No difference in clonality was detected between repertoires from inflamed joints and IFA controls **(figure 3 G)**, but inflamed joints have reduced clonal diversity compared to IFA controls **(figure 3 H)**. This would suggest that antigen driven accumulation of expanded clones in inflamed joints can also be observed at the late phase.

### The antigen experienced endogenous CD4 T cell repertoire displays disparity in clonal diversity between inflamed pLNs and joints with time

Antigen experienced endogenous CD4 T cell repertoires from pLNs and joints were compared between the early and late phases to investigate how CD4 TCR clonality evolves with disease progression **(figures 4 A and E)**. When comparing inflamed pLNs at the early and late phases, the top 10 most expanded clones contributed approximately 18% to the overall antigen experienced repertoire of inflamed pLNs at the early phase compared to only 2% at the late phase **(figures 4 A and B)**. When comparing clonality and diversity, repertoires from pLNs isolated at the late phase displayed increased clonality and diversity compared with their early counterparts **(figures 2 C and D)** and comprised mainly of low abundance clones. In contrast to the pLNs, antigen experienced CD4 T cell repertoires isolated from inflamed joints at the early and late phases were similar; the top 10 most expanded clones dominated the repertoires and comprised on average 46% and 41% of the overall populations **(figures 4 E and F)**. Furthermore, no differences were observed between the clonality and diversity of the repertoires at the early and late phases **(figures 4 G and H)** and repertoire diversity overall was low in the joint at both phases. Together, these data highlight the disparity in repertoire composition between pLNs and joints at the early and late phases, namely that pLNs at the late phase displayed high repertoire clonality and diversity compared to their early counterparts and also compared to inflamed joints at the same phase.

**Figure 4.**
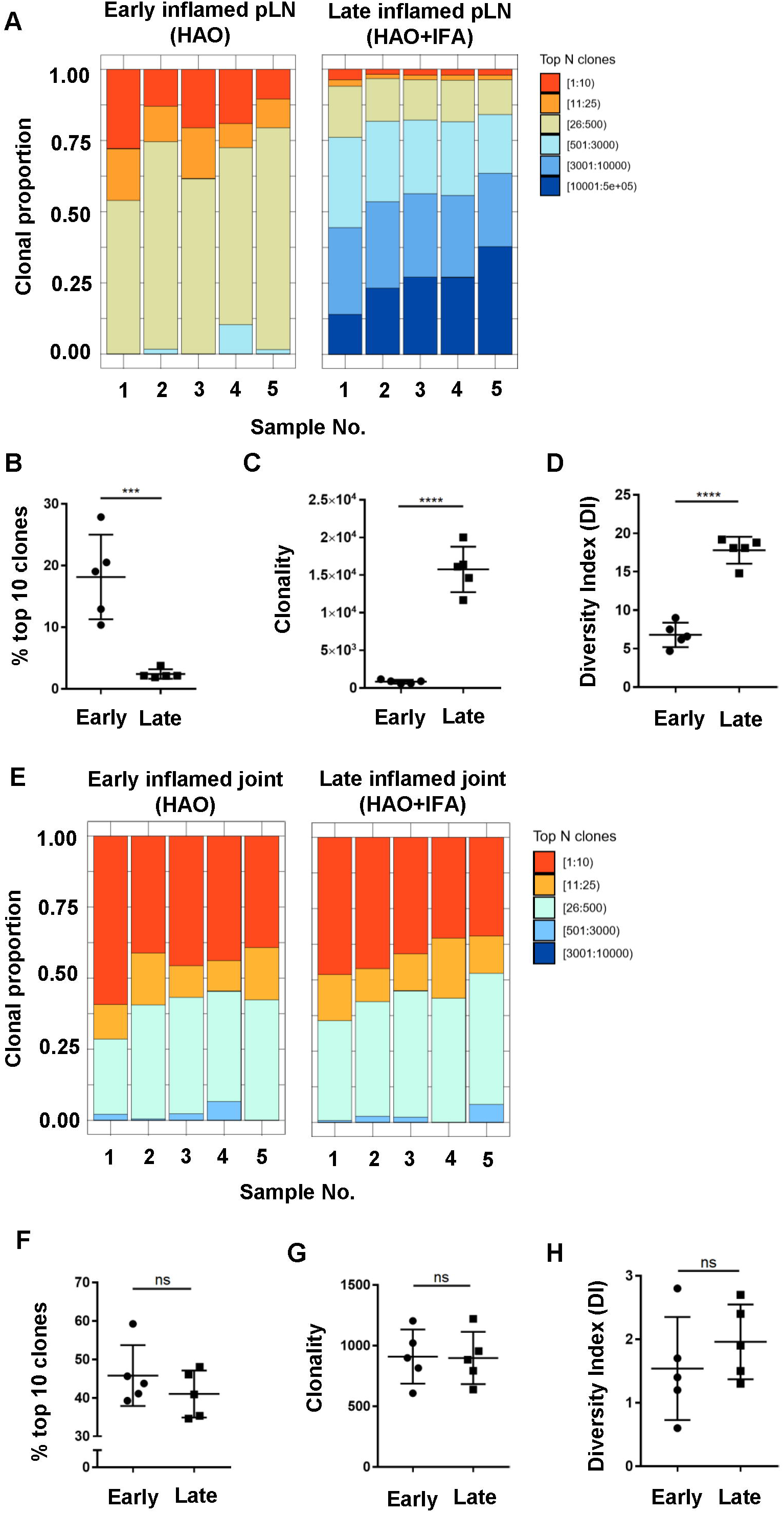
Characterisation of antigen experienced endogenous CD4 T cell repertoires in inflamed pLNs and joints at the early and late phases. Proportion of the top 10 most frequently occurring clones, followed by the next 25, 500, 3000, 10,000, and 100,000 most frequent clones contributing to the endogenous antigen experienced CD4+ T cell repertoire in **A)** inflamed pLNs at the early and late phases, and **E)** inflamed joints at the early and late phases. **B and F** percentage contribution of the top 10 most frequently occurring clones in **B)** inflamed pLNs and **F)** inflamed joints at the early and late phases. **C and G** clonality, represented as the number of unique CDR3β DNA sequences, in **C)** inflamed pLNs, and **G)** inflamed joints at the early and late phases. **D and H** diversity indices of **D)** inflamed pLNs, and **H)** inflamed joints at the early and late phases. Data is representative of mean ± SD with each point representing individual experimental mice. Data represents 1 experiment with n=5 for both the early and late phase groups. Groups were compared using unpaired Student’s t-test. Stars represent the following p values: *** <0.001; **** <0.0001; ns: not significant.

### Disparity in clonality of the antigen experienced endogenous CD4 T cell repertoires is attributed to accumulation of clones with different TCRβ variable (TRBV) genes

Characterisation of the antigen experienced CD4 T cell repertoires from pLNs and joints at the early and late phases provided useful information on the overall distribution of clones in these repertoires. However, this does not provide any insight on the TCR sequence of these clones, nor on whether these clones have similar or different TCR sequences between the two sites. Moreover, no insight is gained on whether the clonal composition of the antigen experienced repertoires in these two sites change over time. To address this, a PCA analysis was performed on inflamed pLNs and joints at both the early and late phases on the basis of TRBV gene use **(figure 5 A)**. Repertoires isolated from inflamed pLNs at the late phase displayed the largest range in TRBV gene use, while inflamed pLNs and joints at the early phase as well as inflamed joints at the late phase grouped together in terms of TRBV gene expression. Indeed, when quantifying these differences, inflamed pLNs at the late phase had significantly increased presence of TRBV genes 1, 2, 3, 5, and 19, while TRBV genes 12-1, 12-2, and 13-1 were present in the highest frequencies in inflamed pLNs at the early phase **(figure 5 B)**. These differences are also due to changes in TRBV gene use and not in TRBJ genes **(figure 5 C)**. Interestingly, the PCA analysis also highlighted inter-sample differences in TRBV gene use in inflamed pLNs at the late phase. Accumulation of clones expressing different TRBV genes in inflamed pLNs at the late phase suggests changes in antigen specific responses with time and also between inflamed pLNs and their associated tissues. Moreover, these changes in antigen specific responses also differ between individual animals with time.

**Figure 5.**
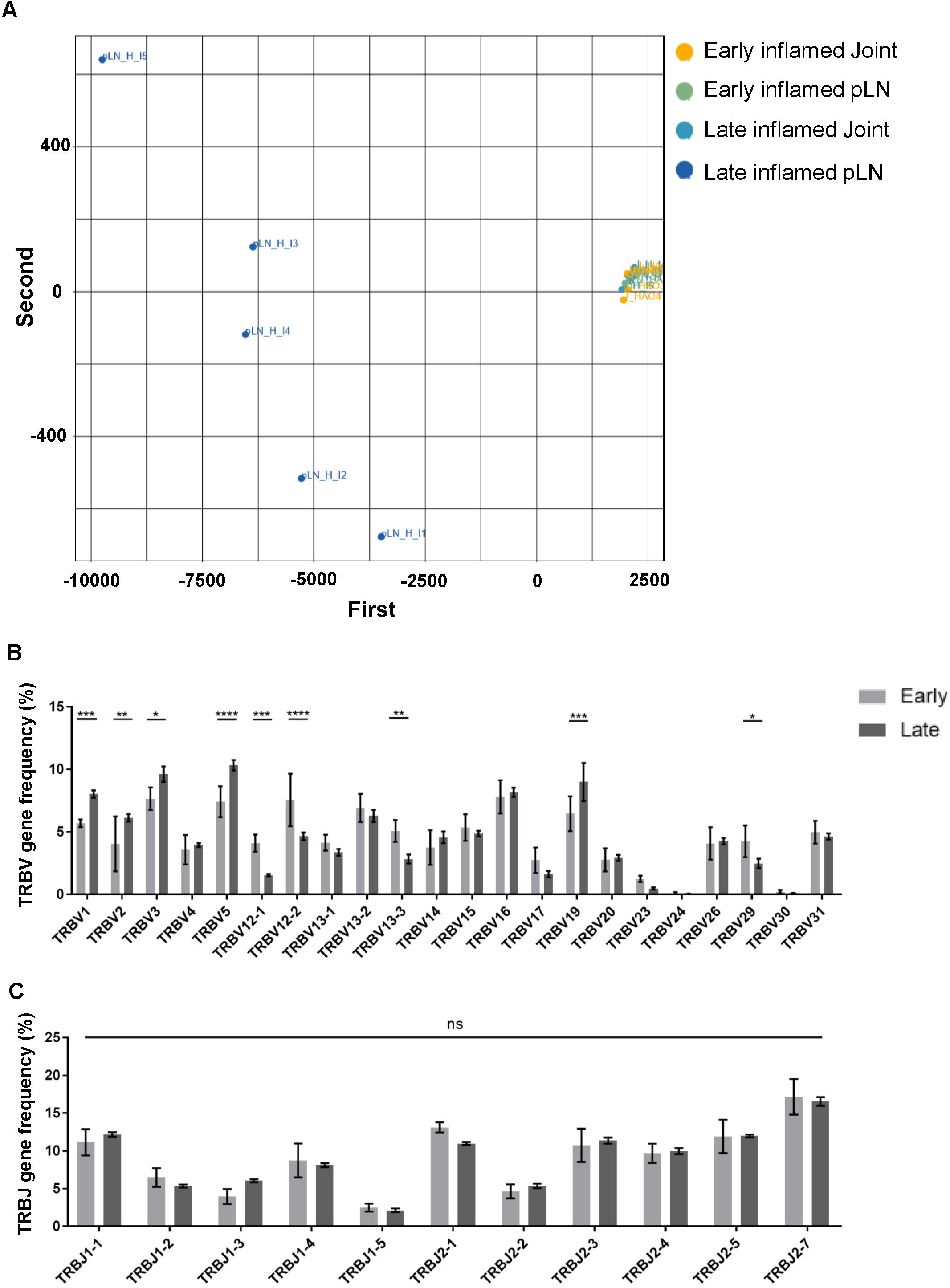
Range of TRBV genes expressed in antigen experienced endogenous CD4 T cell repertoires of inflamed pLNs and joints at the early and late phases. **A)** PCA plot of TRBV gene frequency in repertoires of inflamed pLNs and joints at the early and late phases. Frequency of **B)** TRBV gene expression, and **C)** TRBJ gene expression in inflamed pLNs at the early and late phases. Graphs represent 1 experiment with n=5 for both the early and late inflamed groups. Groups were compared using ordinary 2-way ANOVA with Sidak’s multiple comparisons test. Data presented as mean ±SD. Stars represent the following p values: * <0.05; ** <0.01; *** <0.001; **** <0.0001; ns: not significant.

### Antigen associated changes in the inflamed pLN may predict the changes in antigen specificities and clonality of the CD4 T cell repertoire in inflamed joints

To assess whether changes in CD4 TCR clonality reflected changes antigen specificities, the degree of CDR3β amino acid sequence overlap between the antigen specific CD4 T cell repertoires was investigated. Comparing the repertoire overlap between inflamed pLNs and joints at both the early and late phases allowed us to monitor clonal dynamics between these two sites with the progression of inflammatory arthritis. The number of overlapping CDR3β amino acid sequences is represented by the normalised overlap index (see methods) and this decreased between the inflamed pLN and its joint with the progression of inflammatory arthritis **(figures 6 A and B)**. CDR3β sequence overlap decreased significantly between samples when comparing repertoires from inflamed pLN samples at the early phase to inflamed pLNs at the late phase **(supplementary figure 3 A)**, while the degree of CDR3β amino acid sequence overlap remained unchanged between repertoires from inflamed joints at the early and late phases **(supplementary figure 3 B)**. These observations confirm the observed changes in TRBV gene use and also imply changes in antigen specific responses in pLNs, which progress towards polyclonality with disease progression. Correlation analysis revealed that the top 10 most expanded clones in the inflamed joint at the late phase correlated significantly with the most expanded clones in the inflamed pLNs at the same phase **(figure 6 D)**. Furthermore, the 10 most expanded clones in inflamed joints at the early phase and those from IFA controls did not correlate with the most highly expanded clones in their respective pLNs **(figures 6 C and E)**. Moreover, the 10 most expanded clones in inflamed joints at the early phase were not all present in their respective pLNs **(supplementary table 1)**, but the top 10 expanded clones in inflamed joints at the late phase were all present in inflamed pLNs at the same phase **(supplementary table 2)**. These results indicate that despite pLNs progressing towards polyclonality with the development of disease, the most highly expanded antigen experienced clones accumulate in the inflamed joint. Thus repertoire clonality in the pLN may eventually be mirrored in the inflamed joint with disease progression.

**Figure 6.**
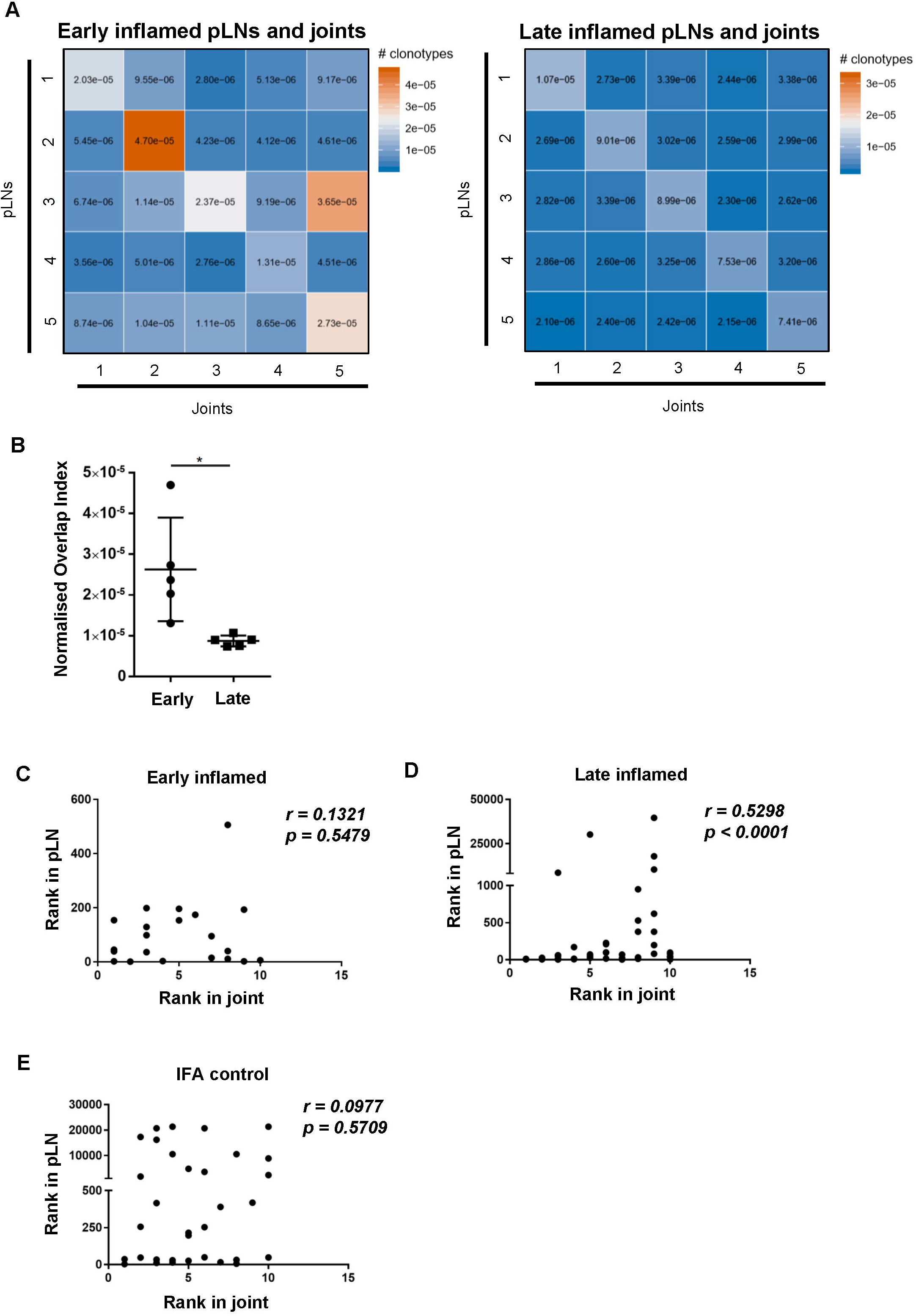
Degree of CDR3β amino acid sequence overlap and correlation of most abundant clones between inflamed pLNs and joints at the early and late phases. **A)** Heatmaps of normalised number of CDR3β amino acid sequences (see methods) between inflamed pLNs and joints at the early (left) and late (right) phases. **B)** Graph representing the intersects (diagonal) of A) representing CDR3β sequence overlap between inflamed pLNs and joint within the same animal. **C-E** Correlation of the top 10 ranked clones in joints with their rank in respective pLN samples. The rank of each of the top 10 clones in **C)** inflamed joint at the early phase, **D)** inflamed joint at the late phase, and **E)** IFA control joint samples were plotted against the rank of that clone in respective pLN samples. Data is representative of one experiment with n=5 for the early and late phase inflamed groups and n=3 for controls. Graph in B) is representative of mean ± SD with each point representing individual experimental mice. * represents p<0.05. In C-E correlations (r) were calculated using Spearman’s correlation after testing for normality of the data using the D’Agostino-Pearson test. p represents the p value.

## DISCUSSION

Characterising CD4 TCR clonality has been used to monitor progression of disease states in cases of infection (32) and autoimmunity (33,34), including RA (15,35–38). Although CD4 T cell clonality has been determined in RA patients at different stages of the disease (15,17), the development of antigen specific responses from very early, pre-clinical stages to a more established disease stage remains unknown. Understanding this is important to develop effective tolerogenic therapies as the range of CD4 T cell clones to target and the timepoint this therapy will have the greatest impact can be determined. Thus, we investigated the evolution of CD4 TCR clonality by sequencing CDR3β regions of antigen experienced endogenous CD4 T cells isolated from pLNs and joints at an early and late phases of breach of self-tolerance models of inflammatory arthritis. Induction of inflammatory arthritis resulted in the accumulation of endogenous CD4 T cells displaying an antigen experienced phenotype. This population has been previously shown to produce TNFα and IFNγ and also harbour autoreactive clones capable of causing bone and cartilage degradation (18–21,39,40). CDR3β analysis of this this population revealed that TCR repertoires from inflamed joints at both the early and late phases displayed low clonality and diversity and were dominated by few clones present at high frequencies. In contrast, pLNs at the late phase displayed higher clonality and diversity compared with their counterparts at the early phase and were mainly composed of a large number of clones that were present at low frequencies. This indicates that pLNs become more polyclonal with the progression of inflammatory arthritis and this polyclonality occurs in the pLNs before possibly being reflected in inflamed joints. This is reinforced by the observations that the most frequently occurring clones in the inflamed joint at the late phase are also the most frequently occurring clones in the inflamed pLN at the same phase. In addition, a study by Klarenbeek et. al (15) demonstrated that repertoires from synovial fluid of patients with established RA have fewer highly expanded clones compared with the synovial fluid taken from patients 6 months after showing clinical symptoms, demonstrating the reduced clonality of CD4 T cell repertoires in the joints of established RA patients. The polyclonality observed in pLNs at the late phase may reflect changes in antigen specific responses due to the release of neo-antigens resulting from continued joint damage and/or epitope spreading, a phenomenon observed in RA patients (41). Indeed, PCA analysis of TRBV genes of repertoires isolated from inflamed pLNs and joints at the early and late phases highlighted the expanded range of TRBV genes present in repertoires of pLNs at the late phase, indicating a possible change in antigen specific responses with the progression of inflammatory arthritis. However, this can also be explained by the re-establishment of lymphatic recirculation. Given that the observations reported were exclusively on the antigen experienced CD4 T cell population, re-establishment of lymphatic recirculation is an unlikely explanation for the changes in the repertoire clonality and diversity observed. Moreover, the 10 most abundant antigen experienced clones correlate significantly between inflamed pLNs and joints at the late phase and not in IFA controls. Inflamed joints were also shown to have reduced diversity in the antigen experienced repertoire compared to joint IFA controls. By taking these observations together, one can deduce that changes in pLN repertoire clonality, diversity, and TRBV genes is in fact antigen driven.

CDR3β amino acid sequence overlap analysis between repertoires isolated from pLNs and joints at early and late phases of inflammatory arthritis highlighted that fewer clones are shared between pLNs and joints at the late phase compared to the early phase. This indicates that antigen specific responses differ between these sites with time, driven by changes in clonality, diversity, and clones with different TRBV genes in the pLN. Moreover, the reduced clonal overlap at the late phase highlight the fact that inflamed pLNs and joints display different antigen specific responses at the same phase. This disparity has important implications for antigen specific therapies as specific pathogenic clones could be potentially targeted in the lymph node before migrating, accumulating, and possibly performing their effector functions in the joint. Moreover, analysis of clonal compositions can also serve as a biomarker to monitor disease progression and assess the therapeutic impact on changes in CD4 T cell clonality. Indeed, changes in cellular compositions in inguinal lymph nodes has been detected in at-risk and early RA patients (42) and treatment with RA therapeutics have been shown to reduce clonal expansion of CD4 T cells (36).

Together, these data outline the evolution of clonal dynamics between inflamed pLNs and their joints with the progression of inflammatory arthritis in that CD4 TCR clonality, begin by being similar between the two sites then change in the pLN with the progression of the disease before potentially being mirrored in the joint in time. These observations have several implications on improving current antigen specific tolerogenic therapies. Firstly, understanding the clonal landscape gives an indication on the range of CD4 T cells that need to be targeted to improven the chances of modulating disease progression. Secondly, targeting expanded CD4 T cell populations early may be more effective therapeutically as the CD4 T cell repertoire is still restricted during the early phases, thus fewer populations would need to be targeted. Lastly, clonality can be a biomarker of disease progression and current therapeutics can be used to ‘reset’ the clonal landscape to one reflecting the earlier phases of the disease, which may enhance effectiveness of tolerogenic therapies and help reinstate self-tolerance and provide long-lasting remission in RA patients.

## ETHICS STATEMENT

This study was carried out at the University of Glasgow and performed in accordance with the UK Home Office regulations and approved by the University of Glasgow Ethics Committee.

## CONFLICT OF INTEREST

The authors declare that the research was conducted in the absence of any commercial or financial relationships that could be construed as a potential conflict of interest.

## ACKNOWLEDGMENTS

We thank the staff within the Institute of Infection, Immunity and Inflammation Flow Cytometry Facility and the Central Research Facility at the University of Glasgow for technical assistance. We also thank Dr. Megan K. MacLeod for her insight and for the critical discussions crucial for the development of this project.

## FUNDING

This work was supported by Versus Arthritis. **RAB, JMB** and **PG** were supported by the Arthritis Research UK (ARUK) programme grant number 19788 and the Innovative Medicines Initiative EU-funded project Be The Cure (BTCURE) [grant number 115142-2]. **JMB** and **PG** were also supported by the Arthritis Research UK Rheumatoid Arthritis Pathogenesis Centre for Excellence (RACE) (grant number 20298) and the Research into Inflammatory Arthritis Centre Versus Arthritis (RACE) (grant number 22072). This work has received support from the EU/EFPIA Innovative Medicines Initiative 2 Joint Undertaking RTCure grant no. 777357. **SAK** was supported by the Ministry of Higher Education Oman (MOHE) throughout the study.

## AUTHOR CONTRIBUTIONS

S.A.K and R.A.B designed the research and performed the experiments. S.A.K analysed data, constructed figures, and wrote the manuscript. C.T.P and J.I.G provided technical assistance and helped with analysis. T.D.O provided technical assistance with the sequencing data, and J.M.B and P.G designed the research and contributed to writing the paper.

---

**Supplementary figure 1**

Gating strategy to identify antigen experienced endogenous CD4 T cells from pLNs and joints. Representative flow cytometry plots identifying antigen experienced endogenous CD4 T cells (CD4+, CD45+, C45.1-, CD44hi) from **A)** inflamed pLNs, and **B)** inflamed joints at the early phase of mice undergoing inflammatory arthritis.

**Supplementary figure 2**

Representative flow cytometry plots identifying antigen experienced endogenous CD4 T from **A)** pLN, and **B)** joint PBS controls at the early phase, **C)** pLN, and **D)** joints IFA controls at the late phase, and **E)** inflamed pLN, and **F)** inflamed joints at the late phase.

**Supplementary figure 3**

Degree of CDR3β amino acid sequence overlap between inflamed pLN and inflamed joint samples at the early and late phases. Heatmaps of normalised number of CDR3β amino acid sequences between **A)** inflamed pLNs, and **C)** inflamed joints at the early and late phases. **B and D** Normalised overlap values between **B)** inflamed pLN samples, and **D)** inflamed joint samples at the early and late phases. Data is representative of mean ± SD with each point representing individual experimental mice. Data represents 1 experiment with n=5 for both the early and late inflamed groups. Groups were compared using unpaired Student’s t-test. Stars represent the following p values: ** <0.01; ns: not significant.

**Supplementary table 1**

The top 10 frequently occurring clones inflamed joint samples at the early phase ranked in terms of frequency in respective pLNs. The top 10 frequently occurring clones in the antigen experienced CD4+ T cell repertoire were found in each of the inflamed joint samples at the early phase and searched for in respective pLN samples. The CDR3β amino acid sequence, associated TRBV gene, read count, and rank of the clone in the pLN sample was obtained. The rank of each of the top 10 clones in the joint are also included in the table. Data is representative of 1 experiment with n=5.

**Supplementary table 2**

The top 10 frequently occurring clones inflamed joint samples at the late phase ranked in terms of frequency in respective pLNs. The top 10 frequently occurring clones in the antigen experienced CD4+ T cell repertoire were found in each of the inflamed joint samples at the late phase and searched for in respective pLN samples. The CDR3β amino acid sequence, associated TRBV gene, read count, and rank of the clone in the pLN sample was obtained. The rank of each of the top 10 clones in the joint are also included in the table. Data is representative of 1 experiment with n=5.

## Notes

https://www.ebi.ac.uk/ena/browser/view/PRJEB40509

### Competing Interest Statement

The authors have declared no competing interest.

## REFERENCES

1. Firestein GS, McInnes IB. Immunopathogenesis of Rheumatoid Arthritis. Immunity (2017) 46:183–196. doi:10.1016/j.immuni.2017.02.006

2. Diaz-Gallo L-M, Martin J. PTPN22 splice forms: a new role in rheumatoid arthritis. Genome Med (2012) 4:13. doi:10.1186/gm312

3. Holoshitz J. The rheumatoid arthritis HLA-DRB1 shared epitope. Curr Opin Rheumatol (2010) 22:293–298. doi:10.1097/BOR.0b013e328336ba63

4. Lee H-S, Lee AT, Criswell LA, Seldin MF, Amos CI, Carulli JP, Navarrete C, Remmers EF, Kastner DL, Plenge RM, et al. Several regions in the major histocompatibility complex confer risk for anti-CCP-antibody positive rheumatoid arthritis, independent of the DRB1 locus. Mol Med Camb Mass (2008) 14:293–300. doi:10.2119/2007-00123.Lee

5. Okada Y, Wu D, Trynka G, Raj T, Terao C, Ikari K, Kochi Y, Ohmura K, Suzuki A, Yoshida S, et al. Genetics of rheumatoid arthritis contributes to biology and drug discovery. Nature (2013) 506:376.

6. Plenge RM, Padyukov L, Remmers EF, Purcell S, Lee AT, Karlson EW, Wolfe F, Kastner DL, Alfredsson L, Altshuler D, et al. Replication of Putative Candidate-Gene Associations with Rheumatoid Arthritis in >4,000 Samples from North America and Sweden: Association of Susceptibility with PTPN22, CTLA4, and PADI4. Am J Hum Genet (2005) 77:1044–1060. doi:10.1086/498651

7. Remmers EF, Plenge RM, Lee AT, Graham RR, Hom G, Behrens TW, de Bakker PIW, Le JM, Lee H-S, Batliwalla F, et al. STAT4 and the Risk of Rheumatoid Arthritis and Systemic Lupus Erythematosus. N Engl J Med (2007) 357:977–986. doi:10.1056/NEJMoa073003

8. Blair HA, Deeks ED. Abatacept: A Review in Rheumatoid Arthritis. Drugs (2017) 77:1221–1233. doi:10.1007/s40265-017-0775-4

9. van Beers JJBC, Schwarte CM, Stammen-Vogelzangs J, Oosterink E, Božič B, Pruijn GJM. The rheumatoid arthritis synovial fluid citrullinome reveals novel citrullinated epitopes in apolipoprotein E, myeloid nuclear differentiation antigen, and β-actin. Arthritis Rheum (2013) 65:69–80. doi:10.1002/art.37720

10. Burkhardt H, Sehnert B, Bockermann R, Engström A, Kalden JR, Holmdahl R. Humoral immune response to citrullinated collagen type II determinants in early rheumatoid arthritis. Eur J Immunol (2005) 35:1643–1652. doi:10.1002/eji.200526000

11. Ioan-Facsinay A, el-Bannoudi H, Scherer HU, van der Woude D, Ménard HA, Lora M, Trouw LA, Huizinga TWJ, Toes REM. Anti-cyclic citrullinated peptide antibodies are a collection of anti-citrullinated protein antibodies and contain overlapping and non-overlapping reactivities. Ann Rheum Dis (2011) 70:188–193. doi:10.1136/ard.2010.131102

12. James EA, Rieck M, Pieper J, Gebe JA, Yue BB, Tatum M, Peda M, Sandin C, Klareskog L, Malmström V, et al. Citrulline-specific Th1 cells are increased in rheumatoid arthritis and their frequency is influenced by disease duration and therapy. Arthritis Rheumatol Hoboken NJ (2014) 66:1712–1722. doi:10.1002/art.38637

13. Law S, Street S, Yu C-H, Capini C, Ramnoruth S, Nel HJ, van Gorp E, Hyde C, Lau K, Pahau H, et al. T-cell autoreactivity to citrullinated autoantigenic peptides in rheumatoid arthritis patients carrying HLA-DRB1 shared epitope alleles. Arthritis Res Ther (2012) 14:R118. doi:10.1186/ar3848

14. Ikeda Y, Masuko K, Nakai Y, Kato T, Hasanuma T, Yoshino SI, Mizushima Y, Nishioka K, Yamamoto K. High frequencies of identical T cell clonotypes in synovial tissues of rheumatoid arthritis patients suggest the occurrence of common antigen-driven immune responses. Arthritis Rheum (1996) 39:446–453. doi:10.1002/art.1780390312

15. Klarenbeek PL, de Hair MJH, Doorenspleet ME, van Schaik BDC, Esveldt REE, van de Sande MGH, Cantaert T, Gerlag DM, Baeten D, van Kampen AHC, et al. Inflamed target tissue provides a specific niche for highly expanded T-cell clones in early human autoimmune disease. Ann Rheum Dis (2012) 71:1088–1093. doi:10.1136/annrheumdis-2011-200612

16. Stamenkovic I, Stegagno M, Wright KA, Krane SM, Amento EP, Colvin RB, Duquesnoy RJ, Kurnick JT. Clonal dominance among T-lymphocyte infiltrates in arthritis. Proc Natl Acad Sci U S A (1988) 85:1179–1183.

17. VanderBorght A, Geusens P, Vandevyver C, Raus J, Stinissen P. Skewed T-cell receptor variable gene usage in the synovium of early and chronic rheumatoid arthritis patients and persistence of clonally expanded T cells in a chronic patient. Rheumatol Oxf Engl (2000) 39:1189–1201.

18. Prendergast CT, Patakas A, Al-Khabouri S, McIntyre CL, McInnes IB, Brewer JM, Garside P, Benson RA. Visualising the interaction of CD4 T cells and DCs in the evolution of inflammatory arthritis. Ann Rheum Dis (2018) 77:579–588. doi:10.1136/annrheumdis-2017-212279

19. Benson RA, Patakas A, Conigliaro P, Rush CM, Garside P, McInnes IB, Brewer JM. Identifying the Cells Breaching Self-Tolerance in Autoimmunity. J Immunol (2010) 184:6378–6385. doi:10.4049/jimmunol.0903951

20. Maffia P, Brewer JM, Gracie JA, Ianaro A, Leung BP, Mitchell PJ, Smith KM, McInnes IB, Garside P. Inducing Experimental Arthritis and Breaking Self-Tolerance to Joint-Specific Antigens with Trackable, Ovalbumin-Specific T Cells. J Immunol (2004) 173:151–156. doi:10.4049/jimmunol.173.1.151

21. Conigliaro P, Benson RA, Patakas A, Kelly SM, Valesini G, Holmdahl R, Brewer JM, McInnes IB, Paul Garside. Characterization of the anticollagen antibody response in a new model of chronic polyarthritis. Arthritis Rheum (2011) 63:2299–2308. doi:10.1002/art.30413

22. Barnden MJ, Allison J, Heath WR, Carbone FR. Defective TCR expression in transgenic mice constructed using cDNA-based alpha- and beta-chain genes under the control of heterologous regulatory elements. Immunol Cell Biol (1998) 76:34–40. doi:10.1046/j.1440-1711.1998.00709.x

23. Bolotin DA, Poslavsky S, Mitrophanov I, Shugay M, Mamedov IZ, Putintseva EV, Chudakov DM. MiXCR: software for comprehensive adaptive immunity profiling. Nat Methods (2015) 12:380.

24. Nazarov VI, Pogorelyy MV, Komech EA, Zvyagin IV, Bolotin DA, Shugay M, Chudakov DM, Lebedev YB, Mamedov IZ. tcR: an R package for T cell receptor repertoire advanced data analysis. BMC Bioinformatics (2015) 16: doi:10.1186/s12859-015-0613-1

25. Katayama CD, Eidelman FJ, Duncan A, Hooshmand F, Hedrick SM. Predicted complementarity determining regions of the T cell antigen receptor determine antigen specificity. EMBO J (1995) 14:927–938.

26. Turner SJ, Doherty PC, McCluskey J, Rossjohn J. Structural determinants of T-cell receptor bias in immunity. Nat Rev Immunol (2006) 6:883–894. doi:10.1038/nri1977

27. Fazou C, Yang H, McMichael AJ, Callan MFC. Epitope specificity of clonally expanded populations of CD8+ T cells found within the joints of patients with inflammatory arthritis. Arthritis Rheum (2001) 44:2038–2045. doi:10.1002/1529-0131(200109)44:9<2038::AID-ART353>3.0.CO;2-1

28. Iannone F, Corrigall VM, Kingsley GH, Panayi GS. Evidence for the continuous recruitment and activation of T cells into the joints of patients with rheumatoid arthritis. Eur J Immunol (1994) 24:2706–2713. doi:10.1002/eji.1830241120

29. Shadidi KR, Aarvak T, Jeansson S, Henriksen JE, Natvig JB, Thompson KM. T-cell responses to viral, bacterial and protozoan antigens in rheumatoid inflammation. Selective migration of T cells to synovial tissue. Rheumatol Oxf Engl (2001) 40:1120–1125. doi:10.1093/rheumatology/40.10.1120

30. Brennan FM, Hayes AL, Ciesielski CJ, Green P, Foxwell BMJ, Feldmann M. Evidence that rheumatoid arthritis synovial T cells are similar to cytokine-activated T cells: involvement of phosphatidylinositol 3-kinase and nuclear factor kappaB pathways in tumor necrosis factor alpha production in rheumatoid arthritis. Arthritis Rheum (2002) 46:31–41. doi:10.1002/1529-0131(200201)46:1<31::AID-ART10029>3.0.CO;2-5

31. Brennan FM, Smith NM, Owen S, Li C, Amjadi P, Green P, Andersson A, Palfreeman AC, Hillyer P, Foey A, et al. Resting CD4+ effector memory T cells are precursors of bystander-activated effectors: a surrogate model of rheumatoid arthritis synovial T-cell function. Arthritis Res Ther (2008) 10:R36. doi:10.1186/ar2390

32. Howson LJ, Napolitani G, Shepherd D, Ghadbane H, Kurupati P, Preciado-Llanes L, Rei M, Dobinson HC, Gibani MM, Teng KWW, et al. MAIT cell clonal expansion and TCR repertoire shaping in human volunteers challenged with Salmonella Paratyphi A. Nat Commun (2018) 9:253. doi:10.1038/s41467-017-02540-x

33. Gomez-Tourino I, Kamra Y, Baptista R, Lorenc A, Peakman M. T cell receptor β-chains display abnormal shortening and repertoire sharing in type 1 diabetes. Nat Commun (2017) 8:1792. doi:10.1038/s41467-017-01925-2

34. Risnes LF, Christophersen A, Dahal-Koirala S, Neumann RS, Sandve GK, Sarna VK, Lundin KEA, Qiao S-W, Sollid LM. Disease-driving CD4+ T cell clonotypes persist for decades in celiac disease. J Clin Invest (2018) 128:2642–2650. doi:10.1172/JCI98819

35. Cantaert T, Brouard S, Thurlings RM, Pallier A, Salinas GF, Braud C, Klarenbeek PL, de Vries N, Zhang Y, Soulillou J-P, et al. Alterations of the synovial T cell repertoire in anti-citrullinated protein antibody-positive rheumatoid arthritis. Arthritis Rheum (2009) 60:1944–1956. doi:10.1002/art.24635

36. Pierer M, Rossol M, Kaltenhäuser S, Arnold S, Häntzschel H, Baerwald C, Wagner U. Clonal expansions in selected TCR BV families of rheumatoid arthritis patients are reduced by treatment with the TNFα inhibitors etanercept and infliximab. Rheumatol Int (2011) 31:1023–1029. doi:10.1007/s00296-010-1402-9

37. Wagner U, Pierer M, Kaltenhäuser S, Wilke B, Seidel W, Arnold S, Häntzschel H. Clonally expanded CD4+CD28null T cells in rheumatoid arthritis use distinct combinations of T cell receptor BV and BJ elements. Eur J Immunol (2003) 33:79– 84. doi:10.1002/immu.200390010

38. Wagner UG, Koetz K, Weyand CM, Goronzy JJ. Perturbation of the T cell repertoire in rheumatoid arthritis. Proc Natl Acad Sci U S A (1998) 95:14447–14452. doi:10.1073/pnas.95.24.14447

39. Nickdel MB, Conigliaro P, Valesini G, Hutchison S, Benson R, Bundick RV, Leishman AJ, McInnes IB, Brewer JM, Garside P. Dissecting the contribution of innate and antigen-specific pathways to the breach of self-tolerance observed in a murine model of arthritis. Ann Rheum Dis (2009) 68:1059–1066. doi:10.1136/ard.2008.089300

40. Platt AM, Gibson VB, Patakas A, Benson RA, Nadler SG, Brewer JM, McInnes IB, Garside P. Abatacept limits breach of self-tolerance in a murine model of arthritis via effects on the generation of T follicular helper cells. J Immunol Baltim Md 1950 (2010) 185:1558–1567. doi:10.4049/jimmunol.1001311

41. Monach PA, Hueber W, Kessler B, Tomooka BH, BenBarak M, Simmons BP, Wright J, Thornhill TS, Monestier M, Ploegh H, et al. A broad screen for targets of immune complexes decorating arthritic joints highlights deposition of nucleosomes in rheumatoid arthritis. Proc Natl Acad Sci (2009) 106:15867. doi:10.1073/pnas.0908032106

42. Rodrìguez-Carrio J, Hähnlein JS, Ramwadhdoebe TH, Semmelink JF, Choi IY, van Lienden KP, Maas M, Gerlag DM, Tak PP, Geijtenbeek TBH, et al. Brief Report: Altered Innate Lymphoid Cell Subsets in Human Lymph Node Biopsy Specimens Obtained During the At-Risk and Earliest Phases of Rheumatoid Arthritis. Arthritis Rheumatol Hoboken NJ (2017) 69:70–76. doi:10.1002/art.39811

