## Supplemental Figure for "TCRβ Sequencing Reveals Spatial and Temporal Evolution of Clonal CD4 T cell Responses in a Breach of Tolerance Model of Inflammatory Arthritis"

### Sup fig 1

#### A) Early Inflamed pLN (HAO)

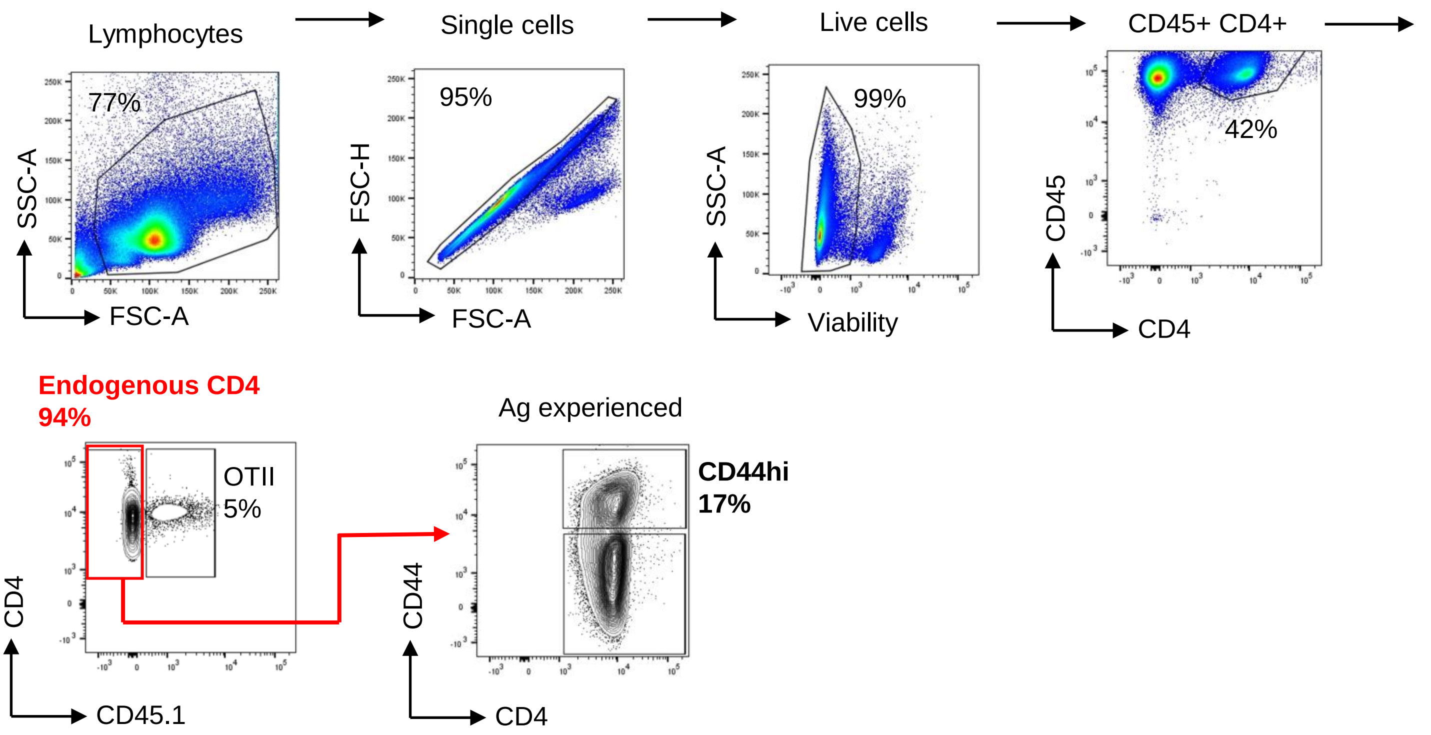


#### B) Early Inflamed Joint (HAO)

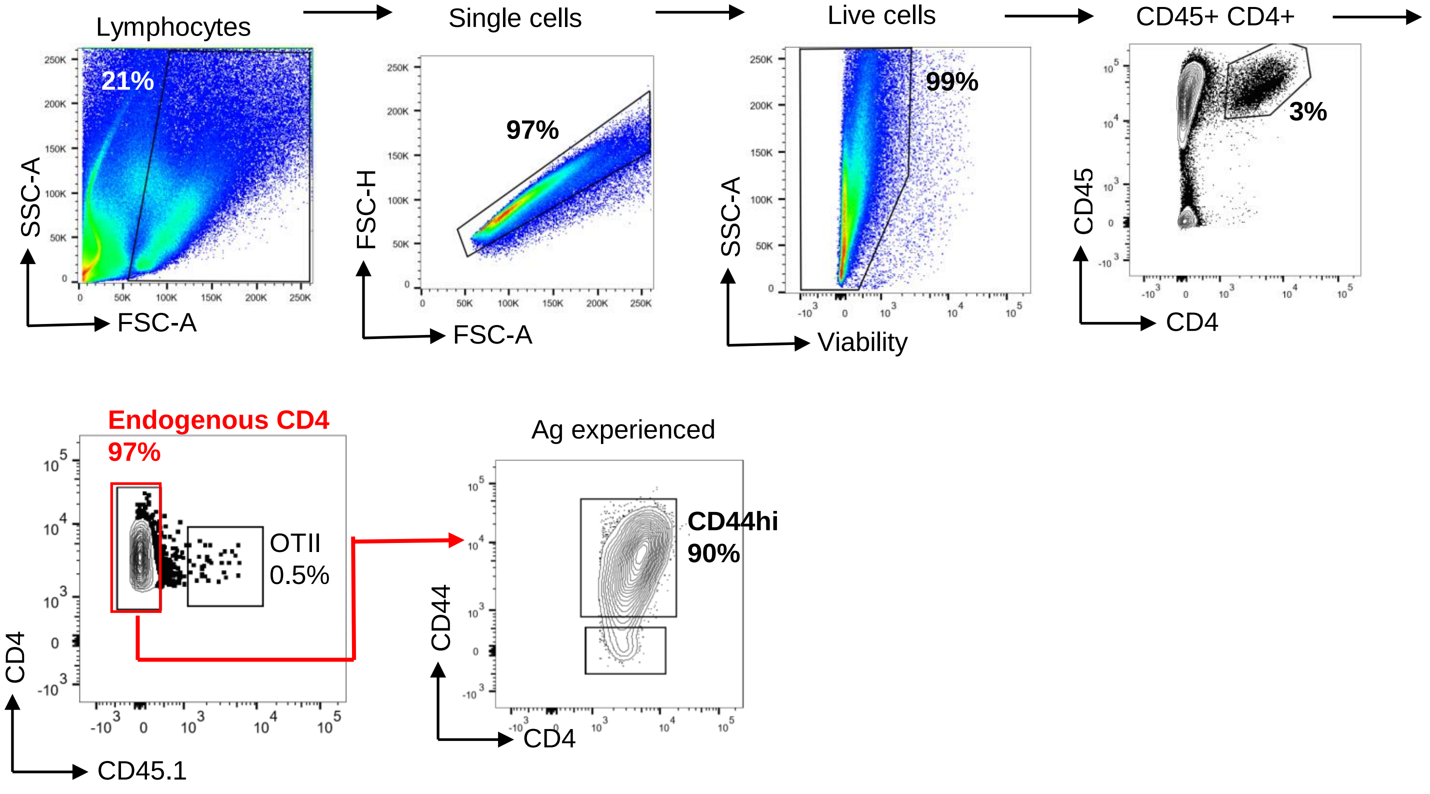

### Sup fig 2

##### A) Early control pLN (PBS)

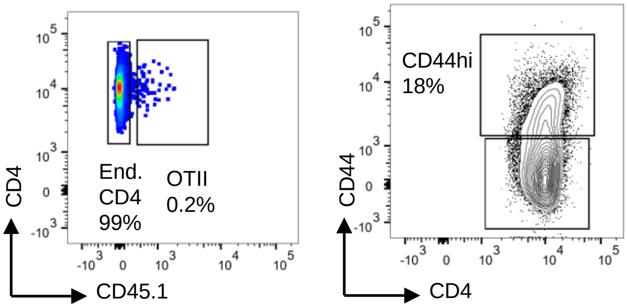


##### B) Early control joint (PBS)

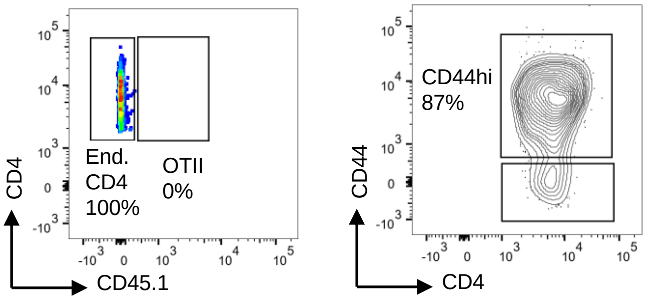


##### C) Late control pLN (IFA)

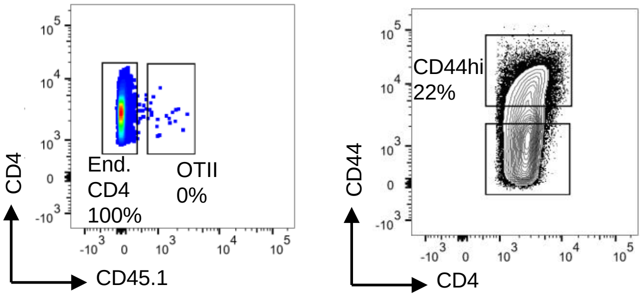


##### D) Late control joint (IFA)

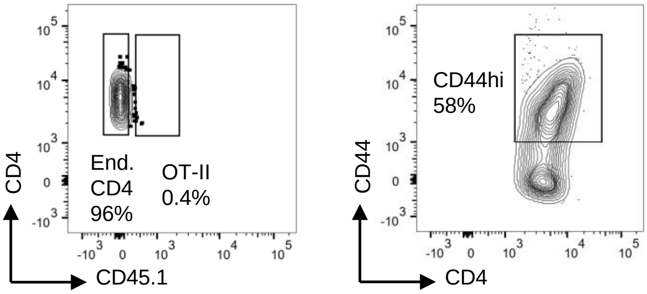


##### E) Late inflamed pLN (HAO+IFA)

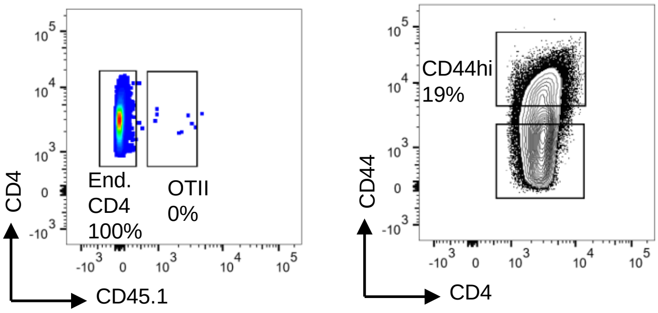


##### F) Late inflamed joint (HAO+IFA)

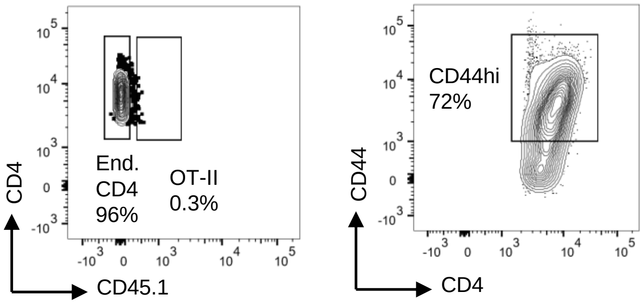

### Sup fig 3

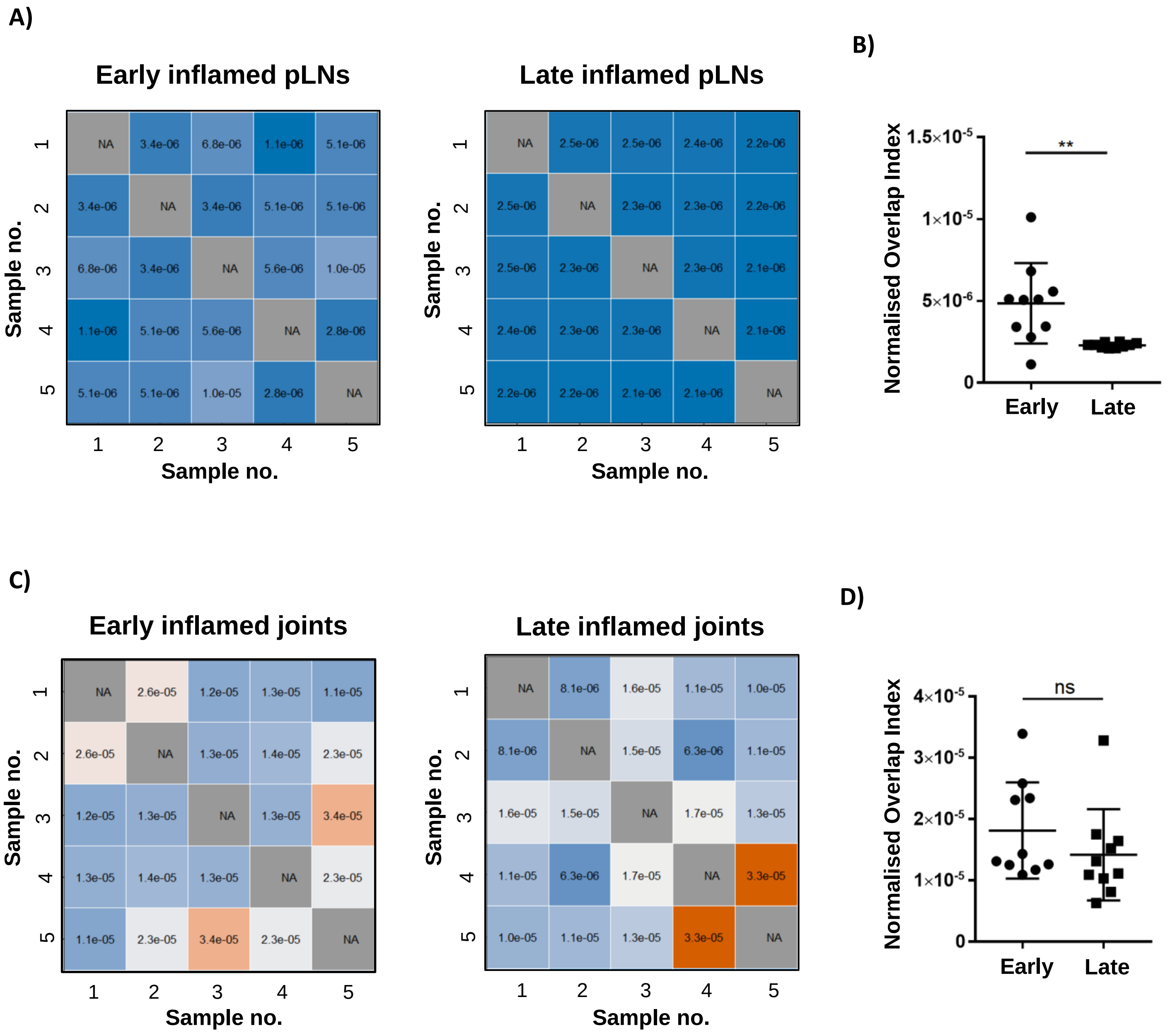
A)
Early inflamed pLNs
Late inflamed pLNs
Sample no.
1 2 3 4 5
NA 3.4e-06 6.8e-06 1.1e-06 5.1e-06
3.4e-06 NA 3.4e-06 5.1e-06 5.1e-06
6.8e-06 3.4e-06 NA 5.6e-06 1.0e-05
1.1e-06 5.1e-06 5.6e-06 NA 2.8e-06
5.1e-06 5.1e-06 1.0e-05 2.8e-06 NA
NA 2.5e-06 2.5e-06 2.4e-06 2.2e-06
2.5e-06 NA 2.3e-06 2.3e-06 2.2e-06
2.5e-06 2.3e-06 NA 2.3e-06 2.1e-06
2.4e-06 2.3e-06 2.3e-06 NA 2.1e-06
2.2e-06 2.2e-06 2.1e-06 2.1e-06 NA
B)
Normalised Overlap Index
1.5×10-5
1×10-5
5×10-6
0
**
Early
Late
C)
Early inflamed joints
Late inflamed joints
NA 2.6e-05 1.2e-05 1.3e-05 1.1e-05
2.6e-05 NA 1.3e-05 1.4e-05 2.3e-05
1.2e-05 1.3e-05 NA 1.3e-05 3.4e-05
1.3e-05 1.4e-05 1.3e-05 NA 2.3e-05
1.1e-05 2.3e-05 3.4e-05 2.3e-05 NA
NA 8.1e-06 1.6e-05 1.1e-05 1.0e-05
8.1e-06 NA 1.5e-05 6.3e-06 1.1e-05
1.6e-05 1.5e-05 NA 1.7e-05 1.3e-05
1.1e-05 6.3e-06 1.7e-05 NA 3.3e-05
1.0e-05 1.1e-05 1.3e-05 3.3e-05 NA
D)
Normalised Overlap Index
4×10-5
3×10-5
2×10-5
1×10-5
0
ns
Early
Late

### Sup table 1

| Sample | CDR3β amino acid sequence | V-gene | Read count in pLN | Rank in pLN | Rank in joint |
|---|---|---|---|---|---|
| Top 10 early joint 1 in early pLN 1 | CASSPGQGTGERLFF | TRBV26 | 4276 | 2 | 1 |
| | CASSQTGGSYEQYF | TRBV2 | 275 | 95 | 7 |
| | CASSNRDWGQDTQYF | TRBV16 | 267 | 98.5 | 3 |
| | CASSFAGHYEQYF | TRBV3 | 94 | 196 | 5 |
| Top 10 early joint 2 in early pLN 2 | CASSLVGEDTQYF | TRBV16 | 1727 | 2 | 9 |
| | CASSPQGNYAEQFF | TRBV2 | 1705 | 3 | 4 |
| | CASSPGGGNYAEQFF | TRBV2 | 1384 | 6 | 10 |
| | CASSLVQNTGQLYF | TRBV16 | 1004 | 15 | 7 |
| | CASSRTGGNSDYTF | TRBV2 | 628 | 39 | 1 |
| | CASSLGAYAEQFF | TRBV3 | 616 | 40 | 8 |
| | CASSKLGGLSYEQYF | TRBV19 | 340 | 98 | 3 |
| Top 10 early joint 3 in early pLN 3 | CASSPLGGRNTGQLYF | TRBV5 | 2307 | 2 | 1 |
| | CASSLGGESQNTLYF | TRBV12-2 | 1250 | 11 | 8 |
| | CASSNREGTQYF | TRBV16 | 753 | 36 | 3 |
| | CGAGQGNTEVFF | TRBV20 | 101 | 193 | 9 |
| Top 10 early joint 4 in early pLN 4 | CASSLGGESQNTLYF | TRBV12-2 | 14841 | 1 | 2 |
| | CASSQGTGGQDTQYF | TRBV2 | 280 | 129 | 3 |
| | CASSFGLEGQNTLYF | TRBV3 | 228 | 154 | 1 |
| | CTCSARGSNTEVFF | TRBV1 | 209 | 174 | 6 |
| | CASSPGQGVNTEVFF | TRBV4 | 61 | 506 | 8 |
| Top 10 early joint 5 in early pLN 5 | CASSQAGGYAEQFF | TRBV2 | 631 | 45 | 1 |
| | CASSQTGVNAEQFF | TRBV2 | 250 | 153.5 | 5 |
| | CASSQNTGQLYF | TRBV16 | 198 | 198.5 | 3 |

### Sup table 2

| Sample | CDR3β amino acid sequence | V-gene | Read count in pLN | Rank in pLN | Rank in joint |
|---|---|---|---|---|---|
| **Top 10 late joint 1 in late pLN 1** | CASSLDLGGRGEQYF | TRBV16 | 976 | 2 | 7 |
| | CASSQTGGAREQYF | TRBV2 | 849 | 3 | 3 |
| | CASSRQGNSDYTF | TRBV12-2 | 807 | 4 | 1 |
| | CASSLSGLGGGAEQFF | TRBV26 | 764 | 5 | 2 |
| | CASGGSGTGETLYF | TRBV13-2 | 456 | 7 | 4 |
| | CASSQAQNTEVFF | TRBV2 | 330 | 11 | 8 |
| | CASSQTGANTEVFF | TRBV2 | 153 | 38 | 10 |
| | CASSLSLGGLEQYF | TRBV26 | 142 | 47 | 5 |
| | CASSSLRGSNQAPLF | TRBV16 | 95 | 98 | 6 |
| **Top 10 late joint 2 in late pLN 2** | CASSFSGQTEVFF | TRBV14 | 530 | 1 | 1 |
| | CASSWGQGRDTQYF | TRBV16 | 267 | 6 | 3 |
| | CASGDLDKYEQYF | TRBV13-2 | 167 | 18 | 2 |
| | CTCSTHSDYTF | TRBV1 | 90 | 69.5 | 7 |
| | CASSQNGALYEQYF | TRBV2 | 89 | 71.5 | 5 |
| | CASSLQSSAETLYF | TRBV16 | 85 | 77.5 | 10 |
| | CASQFRQDNYAEQFF | TRBV14 | 84 | 81 | 9 |
| | CASSPPYINSDYTF | TRBV3 | 60 | 171 | 4 |
| | CASSQAGGYAEQFF | TRBV2 | 52 | 226.5 | 6 |
| | CASSKGPYNSPLYF | TRBV19 | 26 | 950 | 8 |
| **Top 10 late joint 3 in late pLN 3** | CASSLETGGASEQYF | TRBV12-2 | 879 | 1 | 2 |
| | CASRTASGNTLYF | TRBV19 | 798 | 2 | 4 |
| | CARSTRAYNSPLYF | TRBV19 | 257 | 7 | 7 |
| | CASSSNTGQLYF | TRBV16 | 217 | 12 | 1 |
| | CTCSASGFSNERLFF | TRBV1 | 199 | 16 | 6 |
| | CASSQTGGSYEQYF | TRBV2 | 164 | 25 | 5 |
| | CASSQDSGGRAEQFF | TRBV2 | 158 | 31 | 8 |
| | CASGRNYAEQFF | TRBV13-2 | 134 | 39 | 10 |
| | CASSQDTGGLNTLYF | TRBV2 | 107 | 51.5 | 3 |
| | CASSQPGTNQAPLF | TRBV2 | 52 | 197 | 9 |
| **Top 10 late joint 4 in late pLN 4** | CASSKPGYAEQFF | TRBV19 | 435 | 3.5 | 1 |
| | CASSQEAGRNTGQLYF | TRBV5 | 435 | 3.5 | 10 |
| | CASSQTGAETLYF | TRBV5 | 260 | 14 | 4 |
| | CASSRGPYNSPLYF | TRBV19 | 203 | 20 | 7 |
| | CASSQAGGYAEQFF | TRBV2 | 170 | 26.5 | 2 |
| | CASSSNTGQLYF | TRBV16 | 109 | 59.5 | 3 |
| | CASSLAGGGYAEQFF | TRBV3 | 102 | 68 | 5 |
| | CASSFGSSAETLYF | TRBV29 | 55 | 209.5 | 6 |
| | CASSGQGWSTGQLYF | TRBV2 | 35 | 528 | 8 |
| | CASSDRASSYEQYF | TRBV3 | 32 | 621.5 | 9 |
| | *CASSSNTGQLYF* | *TRBV16* | *6* | *8460* | *3* |
| | *CASSLAGGGYAEQFF* | *TRBV3* | *1* | *30109.5* | *5* |
| **Top 10 late joint 5 in late pLN 5** | CASSLAWGGMAEQFF | TRBV16 | 515 | 3 | 4 |
| | CASSHRLANSDYTF | TRBV14 | 496 | 4 | 1 |
| | CTCSRDRGSGNTLYF | TRBV1 | 255 | 13 | 3 |
| | CASSSNQDTQYF | TRBV16 | 217 | 19 | 7 |
| | CASSPLGNYAEQFF | TRBV2 | 196 | 23 | 5 |
| | CASSPPGQNQAPLF | TRBV2 | 76 | 95.5 | 6 |
| | CASSQMTISNERLFF | TRBV2 | 76 | 95.5 | 10 |
| | CASSLDGRSQNTLYF | TRBV16 | 34 | 377 | 8 |
| | CASSQGQQDTQYF | TRBV2 | 34 | 377 | 9 |
| | *CASSQGQQDTQYF* | *TRBV2* | *5* | *10330* | *9* |
| | *CASSQGQQDTQYF* | *TRBV2* | *3* | *17765.5* | *9* |
| | *CASSQGQQDTQYF* | *TRBV2* | *1* | *39595* | *9* |
